# Transfer learning enables prediction of *CYP2D6* haplotype function

**DOI:** 10.1101/684357

**Authors:** Gregory McInnes, Rachel Dalton, Katrin Sangkuhl, Michelle Whirl-Carrillo, Seung-been Lee, Philip S. Tsao, Andrea Gaedigk, Russ B. Altman, Erica L. Woodahl

## Abstract

Cytochrome P450 2D6 (*CYP2D6*) is a highly polymorphic gene whose protein product metabolizes more than 20% of clinically used drugs. Genetic variations in *CYP2D6* are responsible for interindividual heterogeneity in drug response that can lead to drug toxicity and ineffective treatment, making *CYP2D6* one of the most important pharmacogenes. Prediction of CYP2D6 phenotype relies on curation of literature-derived functional studies to assign a functional status to *CYP2D6* haplotypes. As the number of large-scale sequencing efforts grows, new haplotypes continue to be discovered, and assignment of function is challenging to maintain. To address this challenge, we have trained a deep learning model to predict functional status of *CYP2D6* haplotypes, called Hubble.2D6. We find that Hubble.2D6 predicts *CYP2D6* haplotype functional status with 88% accuracy in a held out test set and explains a significant amount of the variability in *in vitro* functional data. Hubble.2D6 may be a useful tool for assigning function to haplotypes with uncurated function, which may be used for screening individuals who are at risk of being poor metabolizers.

## Introduction

Cytochrome P450 family 2, subfamily D, polypeptide 6 (*CYP2D6*), is one of the most important pharmacogenes. The protein, a hepatic enzyme, metabolizes more than 20% of clinically used drugs including antidepressants, antipsychotics, opioids, antiemetics, antiarrhythmics, β-blockers, and cancer chemotherapeutics^1–3^. In the UK Biobank 11% of participants are on a drug with an FDA label indicating actionable or informative *CYP2D6* pharmacogenetics^4^. *CYP2D6* is highly polymorphic, making it challenging to address clinically^5,6^. More than 140 haplotypes comprised of single nucleotide variants (SNVs), insertions and deletions (INDELs), and structural variants (SVs) have been discovered and catalogued in the Pharmacogene Variation Consortium (PharmVar; www.pharmvar.org), many of which are known to alter enzymatic activity and protein expression levels^7,8^. Individuals can be grouped by their CYP2D6 metabolic function and are typically classified into one of four metabolizer (or phenotype) groups: normal (NM), intermediate (IM), poor (PM), and ultrarapid metabolizers (UM). Phenotype frequencies vary widely among global populations with PMs ranging from 0.5% to 5.4%, IMs ranging from 2.8% to 11%, and UMs ranging from 1.4% to 21.2%^9^.

Despite its highly polymorphic nature, *CYP2D6* is one of the most clinically actionable pharmacogenes. A standardized method to translate *CYP2D6* genotype has been recommended by the Clinical Pharmacogenomics Implementation Consortium (CPIC)^10^. Clinical guidelines providing dosage recommendations for different phenotype groups have been published by CPIC for drugs metabolized by CYP2D6, including opioids, selective serotonin reuptake inhibitors, tricyclic antidepressants, the attention-deficit/hyperactivity disorder drug atomoxetine, the estrogen receptor modulator tamoxifen, among others^11–15^ Following the guidelines for these drugs could improve patient outcomes by decreasing adverse effects or increasing efficacy^16^. It has even been suggested that pharmacogenetics-guided opioid therapy could be part of a solution for combating the opioid epidemic^17^. A major insurance company recently announced coverage of *CYP2D6* genotype among other genes for improved selection of antidepressants, marking a major advancement in the incorporation of pharmacogenomics in patient care^18^.

Pharmacogenetic dosing guidelines presume that the clinician has access to the patient’s *CYP2D6* genotype and that the resulting phenotype can be predicted with accuracy. The system used to translate *CYP2D6* genotype into phenotype is known as the activity score (AS) system, which has been widely adopted and is utilized by CPIC^10,19^. *CYP2D6* haplotypes are named using the star allele nomenclature curated by PharmVar, which defines the core variants for each star allele^6–8^. Core variants are typically coding variants, but can also be functionally important variants such as splice junction variants. The AS works by first assigning a value to each star allele (0 for no function alleles, 0.25 or 0.5 for decreased function alleles, and 1 for normal function alleles; gene duplications receive double the value of their single counterpart), then summing the values assigned to each allele. The resulting AS for the person’s pair of haplotypes (or diplotype) is used to determine the CYP2D6 phenotype, which can then be used to inform treatment decisions^10^.

The scores used for a *CYP2D6* haplotype’s contribution to an individual’s AS rely heavily on the manual curation of star allele function through a review of the literature. Most often, *in vitro* experiments and *in vivo* phenotype measures are used to make a determination of star allele function. Even where *in vitro* functional studies exist, however, it can be difficult to assess haplotype function due to variability between expression systems and substrates.

The current reliance on manual curation for scoring of *CYP2D6* haplotypes limits the ultimate utility of CYP2D6 phenotype prediction for pharmacogenetic guidance of treatment decisions. It is estimated that individuals carrying *CYP2D6* haplotypes with an unknown, uncertain, or uncurated function (herein referred to collectively as uncurated) range from 2 to 9%, with a study of the UK Biobank finding that 8% of individuals carry haplotypes of uncurated function^4,20^. Individuals carrying a haplotype with uncurated function cannot be assigned to a distinct metabolizer group using the AS and are instead labeled as “Indeterminate”. Therefore, for these patients, pharmacogenomic-guided therapy for drugs metabolized by CYP2D6 (e.g. CPIC dosing guidelines) cannot be used. In fact, there are currently over 70 star alleles in PharmVar with unknown, uncertain, or uncurated function. Additionally, the extent of the true population level variation in *CYP2D6* is likely far greater than that which can be explained by existing star alleles. In gnomAD, a large aggregate database of genomes, 544 nonsynonymous SNVs and INDELs are identified in *CYP2D6*, and only 98 of those are included in existing star alleles in PharmVar^21^. Further work is required, however, to determine whether these SNVs can truly be ascribed to *CYP2D6* or whether they are misaligned from *CYP2D7*, a highly homologous pseudogene of *CYP2D6*.

With the ever-increasing amount of sequencing data being generated, the number of novel haplotypes of *CYP2D6* keeps increasing making it even more difficult to generate functional data that fulfil the criteria for function assignment recently described by CPIC^22^. The current framework of manual curation of literature in order to assign function to newly discovered star alleles will be challenged to keep up with the rate at which star alleles are discovered, since there will be a lag between the discovery of the star alleles and the generation of functional data for rare alleles.

Methods for predicting variant deleteriousness *in silico* are abundant, but these methods are often developed to be of general purpose genome-wide and are not gene locus specific. Even when gene-specific methods exist, these methods are focused on the prediction of the impact of single variants, rather than the prediction of the impact of a combination of variants. There are existing methods that can predict function for pairs of variants^23^, but *CYP2D6* star alleles can have as many as ten core variants (e.g. *CYP2D6*57*). To predict function of *CYP2D6* haplotypes *in silico*, a purpose-built tool is needed. Tools have been developed to assign *CYP2D6* star alleles to sequence data, but these tools do not predict star allele function^24,25^.

The rise of big data and machine learning, in particular deep learning, has revolutionized computer vision and is being successfully applied to applications in genetics^26^. Deep learning presents an attractive solution for making functional predictions about variation in highly polymorphic genes, like *CYP2D6*. However, deep learning algorithms require massive amounts of data to be used effectively and using deep learning to address all problems in genetics is not straightforward. Many domains in genetics are data limited and therefore challenging to address with complex algorithms. Data scarcity has been addressed in computer vision through the use of transfer learning, where a neural network is pretrained on a large corpus of data, then adapted to a new domain by transferring the coefficients learned by the network to a new network. This is particularly useful in cases where there are spatial or sequential motifs that are shared between the source and target tasks. A neural network can be pretrained using real data from a related task, or simulated data labeled using an existing knowledge source.

Neural networks offer flexibility in the way data is represented. For computer vision tasks, data is represented using three stacked matrices, or channels, representing the RGB channels used to color pixels. Genetic data is represented using a one-hot encoded 4xN matrix, where there are four channels for each of the possible nucleotides and N represents the sequence length. It is possible to include additional information about each position in the sequence by adding additional channels to this input matrix. This flexibility allows us to include variant annotation information, such as whether a variant is in a coding region or whether it is predicted to be deleterious. A functional representation of genetic variants such as this may provide context for observed variants that the neural network can use to reason about newly observed variants.

Here we present a model for predicting function of *CYP2D6* haplotypes, Hubble.2D6. Hubble.2D6 predicts a functional phenotype of *CYP2D6* haplotypes from DNA sequence data using a convolutional neural network (CNN). The model predicts whether a *CYP2D6* haplotype will have normal, decreased, or no function. We validated our model using *in vitro* studies from literature of 48 star alleles, of which 39 had not previously been seen by the model. We generated predictions for 71 *CYP2D6* star alleles that do not yet have a curated function, which may help in increasing the availability of CYP2D6 phenotype prediction. We trained our model using transfer learning and a functional variant representation, and show that these features greatly improve the ability of the network to learn to predict function.

## Results

We trained Hubble.2D6, a deep learning model that predicts star allele functional phenotype from DNA sequence, to classify *CYP2D6* star alleles as normal, decreased, or no function (Fig. 1). Hubble.2D6 was trained to classify function on a set of star alleles for which a functional label has been assigned by curators. We used a training set of 31 star alleles with curated function and a separate set of 25 star alleles with curated function for model validation. Hubble.2D6 correctly predicts the functional phenotype of 100% of the 31 star alleles used for training and 88.0% of the 25 star alleles used as a held-out validation set (Fig. 2a & b). The only misclassifications among samples with curated function are two decreased function alleles in the test set that were predicted to be no function alleles (*CYP2D6*14* & *CYP2D6*72*). Of the 71 star alleles with unknown, uncertain or not yet curated function, 30 were predicted to be normal function, 36 decreased function, and 5 were predicted to be no function, although the true function of these star alleles remains uncurated (Fig. 2c). Predictions for all investigated star alleles, including those with uncurated function, are provided in Supp. Table 1.

**Figure 1.**
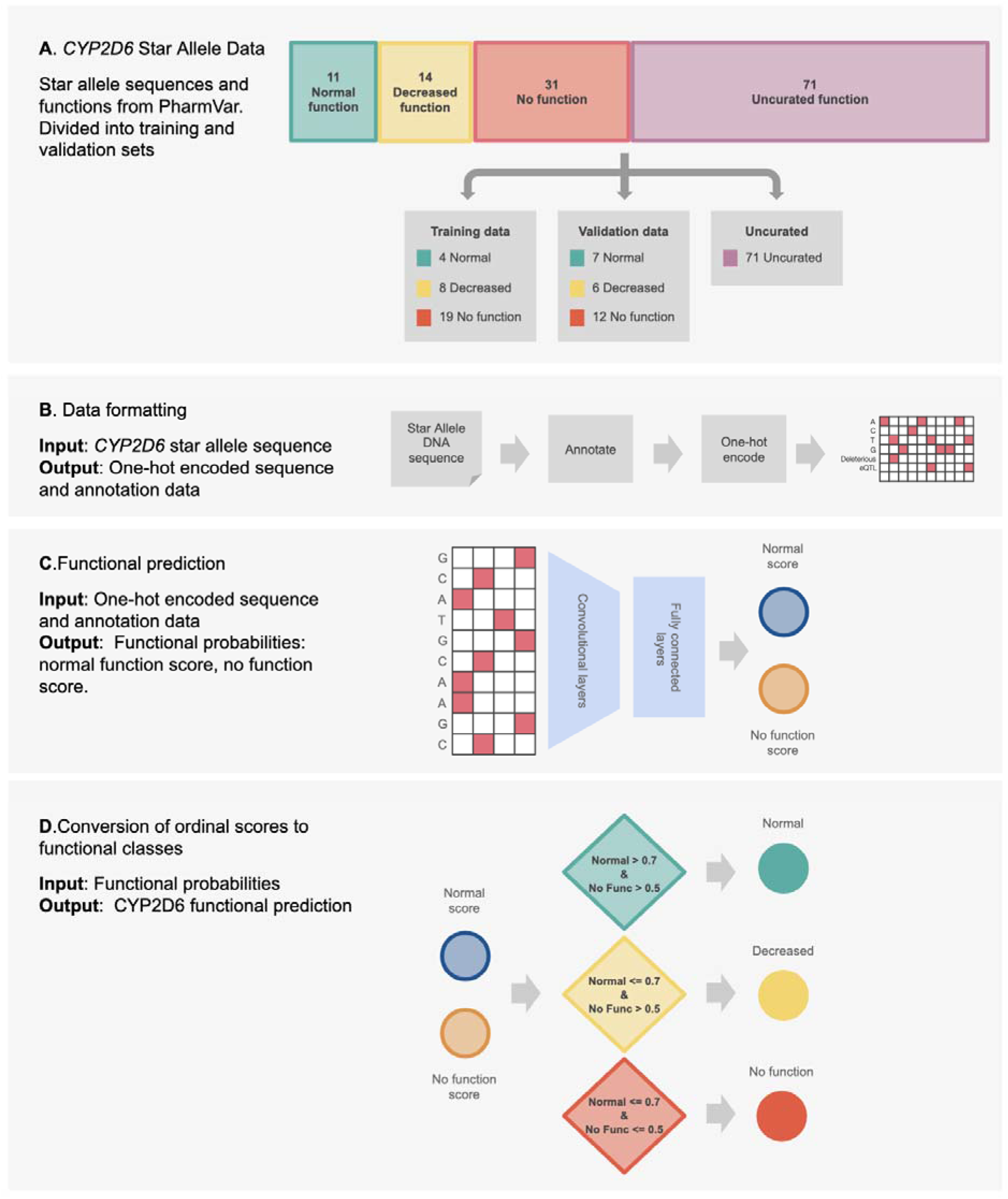
Schematic overview of the Hubble.2D6 workflow. (A) Sequences and functions for all existing star alleles in PharmVar were collected and divided into training and validation datasets. Star alleles with uncurated function were held from training. (B) Star allele sequences were annotated with functional annotations and one-hot encoded as preparation for input into the deep learning model. (C) One-hot encoded sequence and annotation data was read into a convolutional neural network that output scores for two classes: a score indicating a normal functioning allele, and a score indicating a no function allele. (D) The two score outputs from the model were transformed into one of the three functional classes using cutoffs that were set to optimize sensitivity and specificity in the training data.

**Figure 2.**
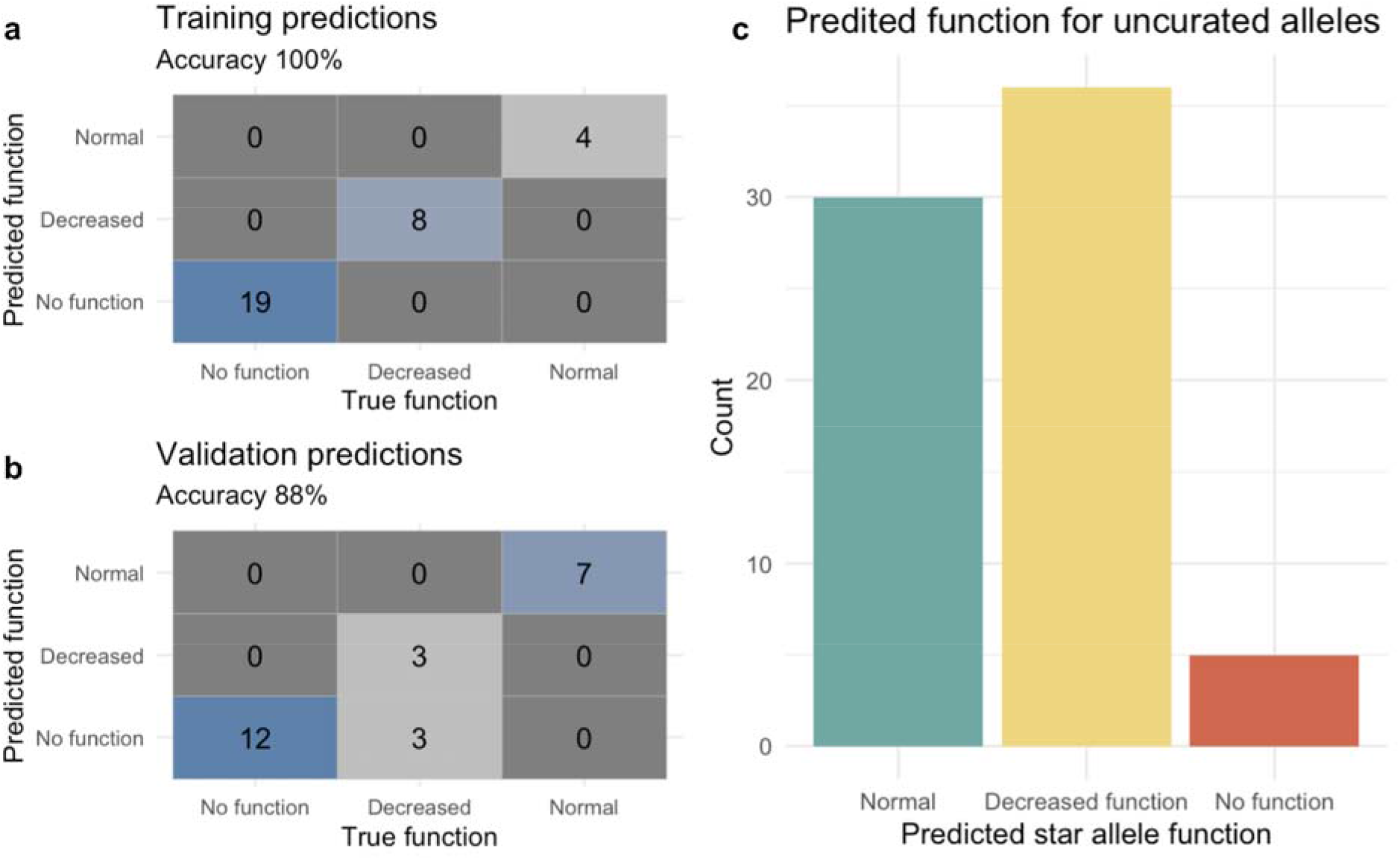
Star allele classification results. The figures depict performance metrics for the prediction of star allele function in the training and validation sets; confusion matrices for class prediction in training and validation are shown in (a) and (b), respectively. (c) shows the frequency of predicted functions for uncurated star alleles.

We evaluated our predictions with *in vitro* data from a study describing functional characterization of 46 *CYP2D6* star alleles^27^. Of the star alleles characterized by this study, 30 have a CPIC assigned functional label. Of the 30 curated function alleles with *in vitro* functional data, 23 were held out from training to be used in the validation set for model evaluation. For these 23 star alleles used for evaluation, our predicted labels explain 71% of the variance, approximately equal to the variance explained by the CPIC assigned labels, 71.1% (Fig. 3a and b). We also assessed the function of 16 star alleles from this study that have not yet been assigned function by CPIC. For these uncurated alleles, two are predicted to have no function, nine decreased function, and five normal function. Our predicted labels explain 47.5% of the variance in the measured activity in the uncurated alleles, the mean measured *in vitro* activity of each predicted phenotype group for the uncurated alleles were significantly different as a result of a one-way ANOVA (*P*=0.014, Fig.3c and d).

**Figure 3.**
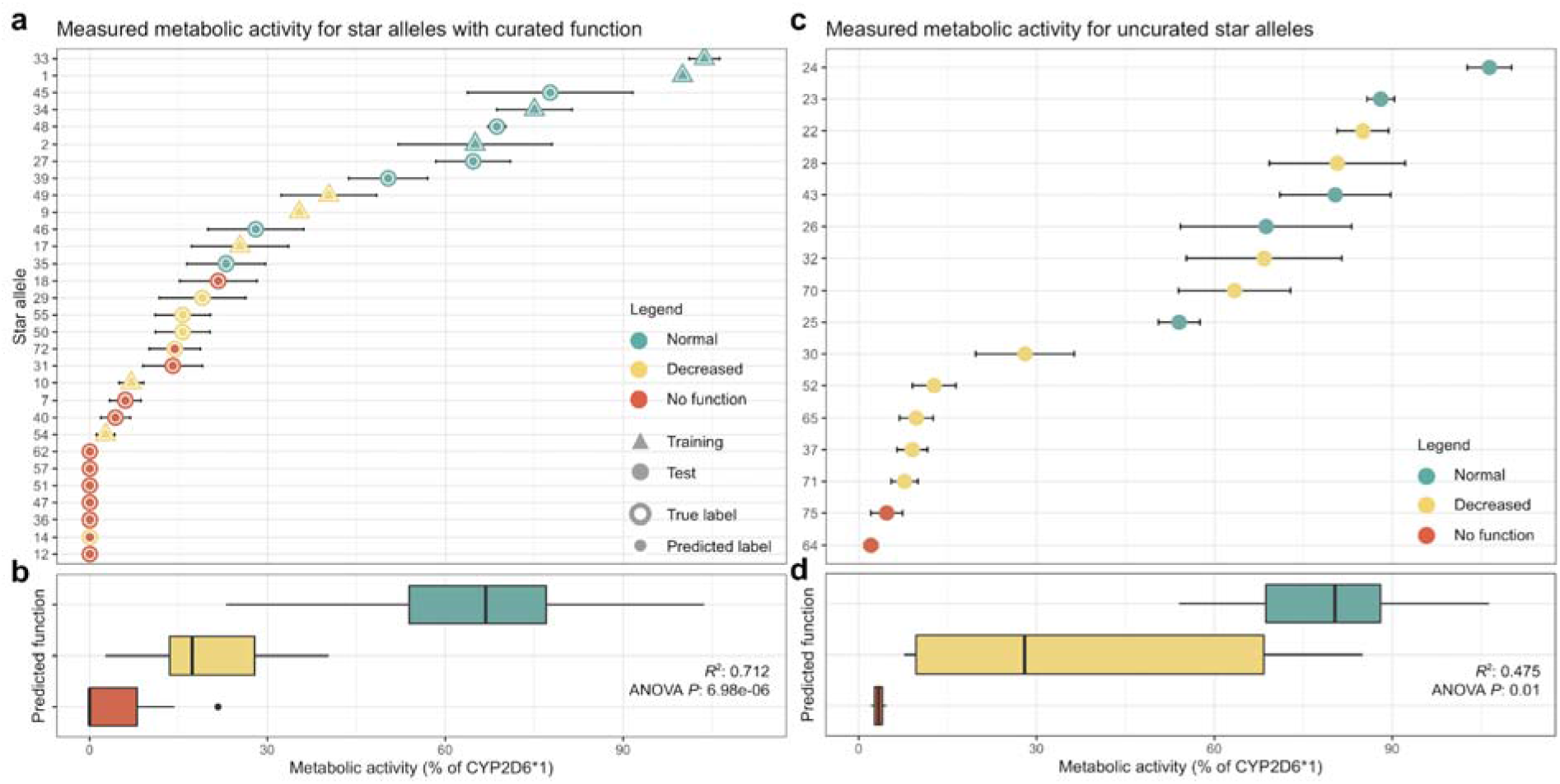
Prediction of star allele function with *in vitro* data. The figures summarize the distribution of metabolic activity measured *in vitro* for star alleles whose function was predicted by Hubble. The distribution of functional activity is shown in A and B for star alleles with curated function. (a) star alleles included in the training process are depicted with a triangle, and those held for testing are depicted with a circle. Error bars depict the standard error of the measured function. The outer edge of each point indicates the true, curator-assigned phenotype, while the inner color represents predicted function. (b) distribution of values for each predicted functional class for data shown in A. (c) star alleles without assigned function status; colors represent the predicted function. (d) variance in measured activity of the star alleles for each predicted label for data shown in C.

We interpreted the predictions made by Hubble.2D6 by calculated importance scores for each variant in each star allele sequence using DeepLIFT. This allows us to see the relative importance of each variant to the final prediction (Fig. 4). Additionally, we wanted to understand whether the model was relying on core variants shared between star alleles to make predictions, or whether novel variants were driving the predictions. We found that, although there is some overlap in core variants between the train and test groups, most star alleles predicted to be of decreased or no function carried unique variants with large importance scores. Importance scores for uncurated star alleles are shown in Supp. Fig. 1.

**Figure 4.**
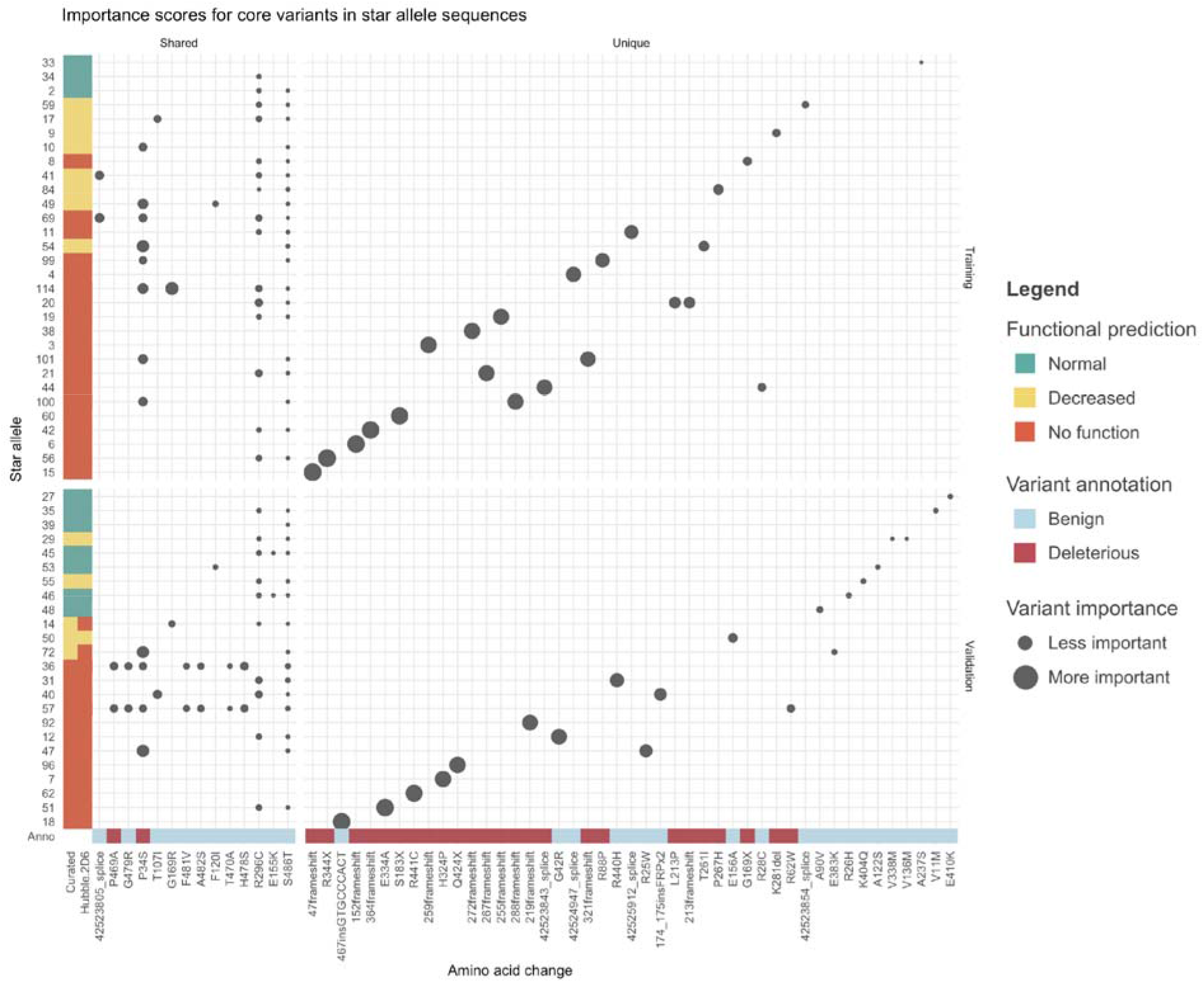
Importance scores for core variants in each star allele used for training and test of Hubble.2D6. Star alleles are along the y-axis and core variants (both amino acid changes and non-coding changes) are listed along the x-axis. Each dot represents the importance of the core variant to the final prediction as determined by DeepLIFT. The size of the dot represents the value of the importance score, with larger dots indicating variants with larger importance scores, typically associated with a negative impact on function. Star alleles are annotated with the curated function as well as the Hubble.2D6 predicted function. Star alleles are divided along the y-axis between star alleles that were included in the training data (top) and those used as test samples (bottom). Star alleles are sorted by the sum of the importance scores, with those with the largest sums at the bottom. Core variants are divided along the x-axis by those that are uniquely in either the training or test samples (right), and those that are shared between star alleles in train and test (left). Core variants are sorted by their mean importance score across all star alleles. Core variants are annotated with the deleteriousness prediction used in the functional variant representation with red indicating a variant predicted to be deleterious and blue indicating a variant predicted to be benign (described in Methods).

We evaluated the contribution of transfer learning and using a functional variant representation by training three new models: two leaving out a single component (one without transfer learning, one without the functional variant representation), and one without either component. We compared the classification accuracy of each of these models to the full model that was trained with both components (Fig. 5). We found that the test accuracy was 28% when excluding both components, 40% for excluding only transfer learning, and 44% excluding only the functional variant annotation, compared to 88% accuracy for the full model. We evaluated each included annotation in the functional variant representation in a similar way. We found that the annotation that most improved the classification accuracy was indicating whether a variant was rare in the population (68% test accuracy), followed by annotating whether a variant is predicted to be deleterious (52% test accuracy).

**Figure 5.**
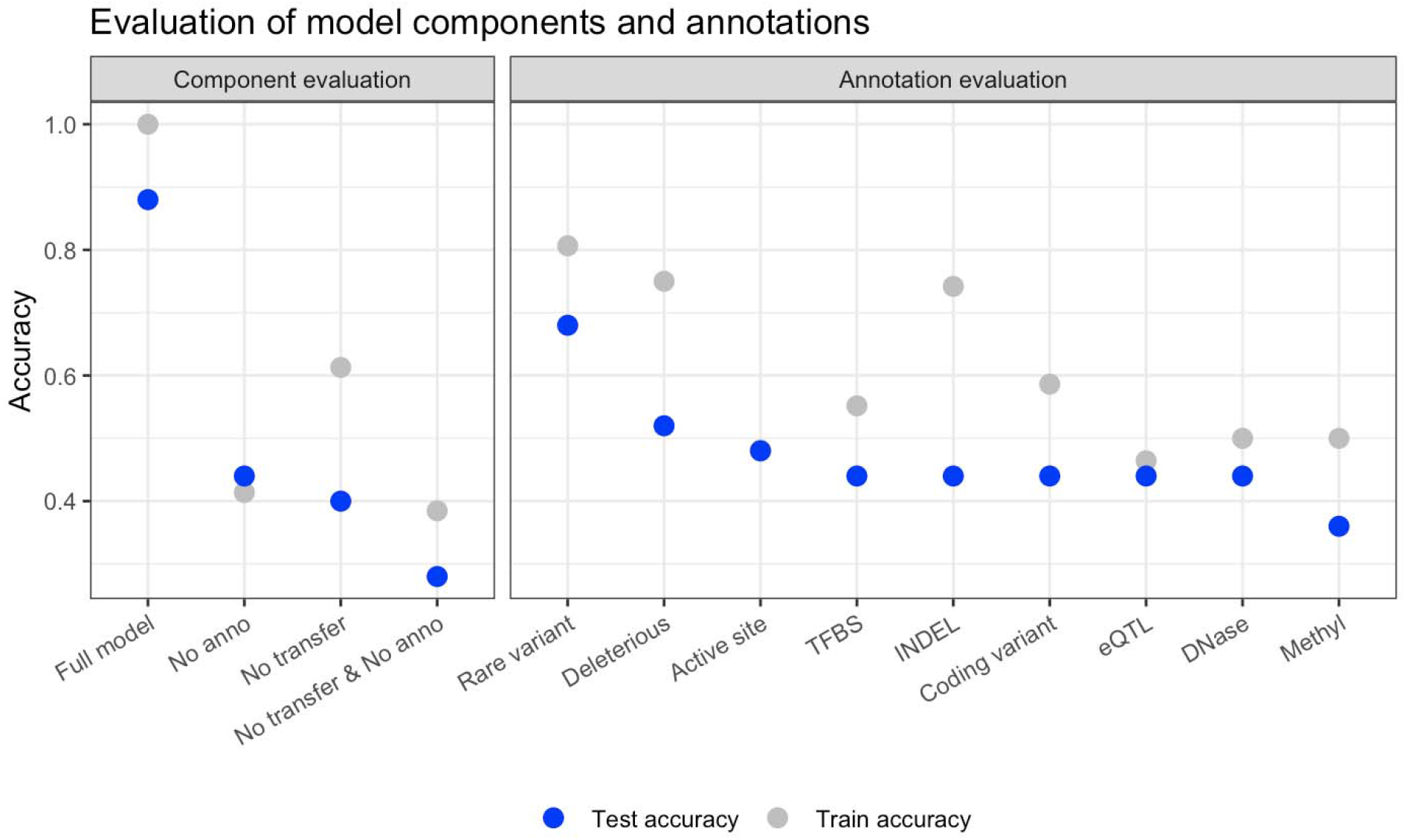
Evaluation of the contribution of deep learning model components. The figure depicts the training and test classification for models trained under various constraints. Under “Component evaluation”, we test the contribution of transfer learning and the inclusion of annotations in the variant encoding. Each were tested individually, together, and one model was built with neither component. Under “Annotation evaluation” we depict in classification accuracy for models trained with a single added annotation. Each point represents the accuracy of a model trained using transfer learning with a one-hot encoding of the nucleotide sequence, but only the specified annotation was included in the encoding of the variant. The full model contains all listed annotations together.

## Discussion

Here we present Hubble.2D6, a model for predicting functional phenotypes of *CYP2D6* star alleles. Given a star allele sequence, Hubble.2D6 classifies the haplotype as a normal, decreased, or no function. Hubble.2D6 has 88.0% accuracy on a held-out validation set of existing haplotypes with curated function. Additionally, we find that in star alleles with uncurated function, the functional predictions from Hubble.2D6 explain a significant amount of the variance in *in vitro* functional measurements, indicating that the Hubble.2D6 assigned labels correlate with actual function.

Predictions of star allele function *in silico*, such as the ones output by Hubble.2D6, can be utilized in several applications in the current pharmacogenomic landscape. Functional predictions could serve as an additional source of evidence when assigning function to star alleles. In cases where *in vitro* data is limited or variable, predicted phenotypes from Hubble.2D6 may be used to guide functional assignment decisions, analogous to a computer-aided decision tool. High throughput assays to generate *in vitro* activity data for large numbers of variants have been suggested for pharmacogenomic applications but have yet to be run comprehensively^28^. Since Hubble.2D6 takes as input the full coding and non-coding sequence of the *CYP2D6* locus, any variant or combination of variants can be rapidly assessed.

Ultimately, functional predictions are anticipated to be part of the clinical prediction of haplotype function in the absence of other data. Currently, if a clinician encounters a patient with a haplotype of uncertain, unknown or uncurated function in a patient, his/her metabolizer status cannot be predicted. Considering the number of star alleles in one of these categories, as many as 8% of patients will not receive a phenotype assignment. Rather than proceed without pharmacogenetic guidance, a phenotype may be predicted using Hubble.2D6. For example, the patient may be more closely followed, or drug level monitoring employed following the initiation of drug therapy, or a drug from a different class selected as a precaution to avoid adverse effects. Such a scenario may be preferable opposed to proceeding blindly without any indication of the patient’s metabolizer status. As we strive to make personalized predictions of drug response a reality for every patient, it is important to be able to provide accurate predicted phenotypes for as many patients as possible.

There are four haplotypes with uncurated function-carrying variants that would likely obliterate function. Star alleles *CYP2D6*81*, *CYP2D6*120*, and *CYP2D6*129* carry nonsense mutations, and *CYP2D6*124* has a frameshift insertion. Of these four alleles, only one was predicted to be a no function allele (*CYP2D6*120*), and the other three were predicted to have decreased function, rather than no function. A prediction of “no function” would be more intuitive, as these types of variants are well known to lead to a non-functional protein. The model importance scores show that the model heavily weighted the variant leading to a premature stop in *CYP2D6*120*, however, the presumably loss-of-function variants in the other star alleles received considerably lower importance scores (Supp. Fig. 1). Each of the four loss-of-function variants is predicted to be deleterious by LOFTEE^29^. It seems that the model treats newly seen variants conservatively and outputs a prediction of decreased function rather than no function. Although this is an important distinction, providing an indication that the allele will metabolize drugs abnormally may still be beneficial. More training data with more varied examples of loss-of-function variants may improve future versions of the model.

An important contribution of this work is the finding that deep learning models can be trained for small-scale problems in genetics. The highly polymorphic nature of *CYP2D6* makes it an attractive application for deep learning, but the limited number of haplotypes with known function to use as training data limits our ability to train large models effectively. We show that through the use of both transfer learning and the inclusion of variant annotations we can train a model to predict haplotype function with high accuracy. The intuition that motivates this is that, first, by pretraining the model on simulated and real data it has learned weights for motifs related to commonly observed variants. This prevents the model from having to learn all motifs from scratch during the training phase. Second, by including variant annotations in the variant encoding the model has been provided with additional useful information. With enough data a model may be able to learn that a certain nucleotide change would be likely to be deleterious, but in a space where data is scarce it may be better to provide the model with that information upfront. As we can see from the variant importance scores in Fig. 4 many of the variants with large importance scores are unique to a single star allele, so it is important for the model to learn the principles that lead to a functional change rather than nucleotide motifs alone. We show that by combining these two methods the result is a more accurate predictor than either method alone.

There are several limitations to our *CYP2D6* predictive models. First, we do not include structural variation in the *CYP2D6* locus in our model. This prevents the model from being able to predict function of star alleles with structural variation including hybrid genes^30^ (e.g. *CYP2D7-2D6* or *CYP2D6-2D7*), gene deletions^31^ (e.g. *CYP2D6*5*), and copy number variants. Thus, Hubble.2D6 does not predict increased function at this point in time. While these are important types of alleles to consider, all known occurrences of hybrid genes result in a complete loss of function, so *in silico* prediction of function is not necessary, and changes in copy number are easily accounted for with the use of the activity score. In the current star allele definitions, the only source of increased function alleles is due to gene duplication events. Second, since we only considered a narrow range of upstream and downstream sequences of *CYP2D6* we have not captured distal effects on gene expression, such as a long-distance ‘enhancer’ SNV that has been described to be associated with decreased function^32–34^. Distal regulatory effects are not included by the current model, which may limit our ability to fully explain the observed variability in enzymatic activity using *CYP2D6* sequence alone. However, these distal effects would not be captured by conventional *in vitro* expression studies either. Third, the validation performed used *in vitro* data for only 50 star alleles, and the data was generated using one expression system in a single laboratory. Substantial variance exists in measured functional activity of star alleles between laboratories. Further functional assessment of star alleles compared to Hubble.2D6 predictions could inspire greater confidence in our ability to predict function *in silico*. Finally, there may be factors impacting CYP2D6 activity outside the gene sequence, such as variation in other genes or whether an individual is taking CYP2D6 inhibitors such as selective serotonin reuptake inhibitors^35,36^. These factors are not included in our model as we only focus on the genetic variation in the *CYP2D6* locus.

In summary, we have created a model for the prediction of metabolic activity for *CYP2D6* haplotypes from sequence data. We find that our model has high accuracy predicting allelic function, and that our predicted function labels explain a significant amount of the variance observed in CYP2D6 metabolic activity *in vitro*. This model may be useful to predict phenotype of patients carrying haplotypes, which are either uncurated or have unknown/uncertain function assignments

## Methods

### Model Overview

We trained a multiclass convolutional neural network (CNN) classifier to predict *CYP2D6* haplotype function from DNA sequence. The deep learning model used is a three layer CNN following the Basset architecture^37^. The output is one of three functional statuses: “No function”, “Decreased function”, or “Normal function”, following the clinical function assignments provided by CPIC and posted by PharmVar, excluding “Increased function” (Fig. 1). Hubble.2D6 does not make predictions for increased function alleles because the only increased function alleles identified for *CYP2D6* occur as a result of gene duplications; only SNVs and small INDELs are considered. Hubble.2D6 reads in a one-hot encoded matrix representing the full DNA sequence of the haplotype in addition to eight variant level functional annotations and outputs two scores: (1) the probability that the haplotype is a no function allele and (2) the probability that the haplotype is a normal function haplotype. The two scores are then transformed into one of the three functional classes (no function, decreased function, or normal function) using cutoffs that are defined to maximize sensitivity and specificity of the functional predictions in the training data. For example, a haplotype receiving a no function probability greater than 0.5 and a normal function probability less than 0.7 is classified as a decreased allele.

Although only two scores are output they are combined to map the function to one of the three possible outcomes; this setup formats the prediction task as an ordinal regression problem such that the network can learn the ranking of the functional groups^38^. The first score, the no function score, differentiates between no function alleles and alleles with either decreased or normal function. No function alleles are indicated with a 0 as the first score while alleles with any other function are indicated with a 1. Likewise, the second score, the normal score, differentiates between alleles with normal function and alleles with either decreased or no function. Normal function alleles have a score of 1 for the second score while others have a score of 0. Leading to a scoring system where each of the three function classes can be yielded from only two scores. This scoring system is superior to a classification setup that assumes class independence because it allows the network to learn that no function alleles are more similar to decreased function alleles than they are to normal function alleles, and vice versa.

The input into the model is genetic sequence data. For each star allele the data is converted into a one-hot encoded matrix of DNA variation and variant-specific functional annotations for each position in the *CYP2D6* locus (chromosome 22, 42,521,567-42,528,984, hg19). The interrogated region includes 2,103 bp of upstream and 934 bp of downstream sequence which corresponds to the region covered by the PGRNseq platform^39^. We used eight variant level annotations that may influence gene expression or protein function to create a functional variant representation. Each base in the capture window was annotated with a binarized annotation for the following characteristics:

1. If the variant is in a coding region, as defined by the RefSeq (NG_047021.1)^40^
2. If it is rare in the population. Defined as allele frequency among all populations in gnomAD < 0.05.
3. If it is deleterious. If it is a coding variant, we use an ADME optimized framework for predicting deleteriousness^41^. If it is non-coding we use a majority vote of CADD, DANN, and FATHMM^41–44^. LOFTEE predictions of deleteriousness supersede the other methods, if available^29^.
4. If it is an INDEL of any length. INDELs are reduced to the first nucleotide and given the INDEL annotation, so as to keep the length of each sequence the same.
5. If it is in a methylation mark, as defined by UCSC Genome Browser tracks wgEncodeHaibMethyl450Gm12878SitesRep1 and wgEncodeHaibMethylRrbsGm12878HaibSitesRep1^45,46^.
6. If it is in a DNase hypersensitivity site. We use UCSC Genome Browser track wgEncodeAwgDnaseMasterSites.
7. If it is in a transcription factor binding site. We use UCSC Genome Browser tracks tfbsConsSites and wgEncodeRegTfbsClusteredV3.
8. If it is a known *CYP2D6* expression quantitative trait loci (eQTL) for any tissue in gTEX v6^47^.
9. If it codes for a residue in the CYP2D6 active site where the substrate binds to the protein^48^.

Variants are annotated using a custom pipeline that includes annovar, VEP, and a script for binarizing the annotations^49,50^. The final dimensions of the input haplotype matrix are 7417×12 (the length of the locus window x the nucleotide vector). Every base in the sequence window, coding and non-coding, is annotated and converted into a 1×12 vector (four possible nucleotides, eight annotations).

### Training procedure

The number of existing star alleles with curated function was small for typical deep learning applications, thus transfer learning was utilized to reduce the amount of training data required to create a robust model. Hubble.2D6 was trained in a stepwise process with two pre-training steps, first with simulated data and second with real data. Each step in the training procedure used a CNN with identical number of convolutional layers and filter shapes, although the model output varied at each step. This allowed the weights of the convolutional layers learned at each step to be transferred to the next stage in training, iteratively updating the weights with different datasets.

First, a CNN classifier was trained to predict the activity score of 50,000 simulated pairs of haplotypes (or diplotypes), thereby creating a neural network representation of the activity score (the simulation procedure is described in the following section). We simulate diplotypes rather than haplotypes in order to be able to further train the model using functional activity data collected from liver microsomes. In the second stage of training, a regression model was trained to predict measured metabolic activity of 314 liver microsome samples using dextromethorphan as substrate^51^. The weights derived from the convolutional layers of the first model were transferred to the second model and the fully connected portion of the model was retrained using randomly initialized weights. The input to this second model was the pair of haplotypes identified through sequencing of the *CYP2D6* loci encoded in the same manner described previously (these data are described in the following section). The final model was created by removing the final output layer of the network trained on the liver microsome data and adding a new output layer with two neurons with sigmoid activations, corresponding to the two outputs described previously (one representing no function status, another representing normal function status). The new network was created by creating a new neural network with an identical architecture, except for the last layer. Then, the weights from the pretrained network are copied to the new network for each layer, except for the final output layer. The final layer with the two output neurons will then be trained from randomly initialized weights in a final training phase using the star allele haplotype sequences. Since the liver microsome model was trained on pairs of haplotypes, the final model input consisted of two identical copies of the input haplotype matrix for star allele classification. The network was then retrained to classify star allele functional status with the starting weights initialized from the liver microsome model. Ten models were trained and their outputs averaged to form an ensemble.

### Training Data

#### Simulations

Simulated diplotypes used in pre-training were created by randomly selecting a pair of *CYP2D6* star alleles with curated function (normal, decreased, or no function haplotypes) that do not have any structural variants and constructing haplotypes with the variants associated with the star alleles. Star allele definitions were downloaded from PharmVar (v4.1.1). To introduce additional diversity to the training data, alternate alleles (both SNVs and INDELs) were sampled for variant sites not associated with any star allele following a uniform distribution with the probability of an alternate allele occurring equal to the population level alternate allele frequency published in gnomAD^21^. It is possible that rare, deleterious variants not currently represented in any star allele were added during this process. However, noisy pretraining data has been shown not to negatively impact the final model^52^. A total of 10,000 genotypes were selected for each AS (0, 0.5, 1, 1.5, 2), for a total of 50,000 simulated samples used in training, and an additional 10,000 total genotypes to use as a test set.

#### Liver Microsome Data

Activity data were available for 314 liver microsome samples from a prior study as a second pretraining step for Hubble.2D6^51^. The liver microsome data was collected from two sites, 249 samples from St. Jude’s Children’s Research Hospital (SJCRH), and 65 samples from University of Washington (UW). All samples were sequenced with the PGRNseq panel^39^. Metabolic activity was measured using two substrates, dextromethorphan and metoprolol, but only dextromethorphan activity was used for pre-training. Star alleles and structural variants were called for each sample using Stargazer^24^.

#### Star allele data

The sequences used to train the final model were constructed based on star allele definitions by PharmVar (version 4.11.1, downloaded 10/25/2019). We selected 31 star alleles (as per the core allele definitions) and each of their suballeles for training. Suballeles are versions of star alleles that have been identified that carry additional variants (typically non-coding variants) other than the core variants that define the star allele (e.g. *CYP2D6*1.002*). The model was evaluated with 24 randomly selected star alleles. Star alleles with no curated function (defined by CPIC and posted by PharmVar as “Uncertain”, “Unknown”, or “Awaits curation”) were excluded from the training procedure. During the training of each model 10% of the samples from each functional class (no function, decreased function, and normal function) were randomly held out to be used to check for overfitting in the training process.

### Validation

The model was evaluated by predicting the function of the 25 star alleles with curated function that were excluded from the training process. The area under the receiver operator characteristic curve (AUROC) was calculated for the training and test groups for the two scores output by the model. In addition, the function of 71 star alleles with uncurated function was predicted.

In order to further validate our model, we used *in vitro* data from a study^27^ that measured the activity of 46 star alleles (including wild-type) using three substrates. Of these 46 star alleles, 30 have curated function while 16 do not have a CPIC assigned function. The metabolic activity used for each star allele was the percent activity of the reference, taking the mean activity across all three substrates. We calculated the variance explained by the predicted function using a linear model and assessed the heterogeneity in the measured activity of each functional group (no function, decreased function, and normal function) using a one-way ANOVA. This was done separately for samples with curated function and those with uncurated function. *CYP2D6*53* was excluded from this analysis because it had a 28-fold increase in *in vitro* function compared to reference, and deemed to be an extreme outlier. Additionally, *CYP2D6*61* and *CYP2D6*63* were excluded because they are *CYP2D6-CYP2D7* hybrids, a result of structural variation which is not included in the Hubble.2D6 framework.

### Model Interpretation

We interpreted the model and the relative importance of the variants in each star allele for the predicted function using DeepLIFT from the DeepExplain package^53,54^. DeepLIFT compares the activations of a neural network for a given sample against a reference sample, and outputs importance scores for each input feature. We ran DeepLIFT on each star allele sequence with a *CYP2D6*1* reference sequence. This yielded importance scores for each variant in each star allele that were different from the variants in *CYP2D6*1*.

### Model Evaluation

We evaluated the added components of the model (transfer learning and the functional variant representation) by training new models with each component removed. Concretely, to evaluate the contribution of transfer learning to the final model, we trained a new model identical to the final model except the weights were randomly initialized rather than transferred from a pretrained network. To evaluate the contribution of the functional variant representation we trained a new model and input only the one-hot encoding of the nucleotide sequence, no annotations included. For this model weights were transferred from a pretrained network. Finally, we trained a third model with that did not include any variant annotations and had randomly initialized weights. For each test case, we followed the same procedure to generate predictions as with the full model: an ensemble of seven models was trained and the average score for each of the two output scores taken across all seven models which was then converted into a single functional class. Classification accuracy was calculated for star alleles in training and test.

We also evaluated each of the annotations included in the functional variant representation. This was done by again training ensemble models with transfer learning, but in this case a single annotation was included in addition to the one-hot encoding of the nucleotide sequence. This was done for each of the eight annotations included in the final model. Again, the mean scores from each ensemble were taken and converted into predictions as previously described and then calculated classification accuracy.

## Acknowledgements

G.M. is supported by the Big Data to Knowledge (BD2K) from the National Institutes of Health (T32 LM012409). E.L.W. and R.D. are supported by the Northwest Alaska-Pharmacogenomics Research Network (NWA-PGRN) (P01GM116691). K.S., M.W.C., and R.B.A are supported by NIH/National Institute of General Medical Sciences PharmGKB resource, (R24GM61374). R.B.A. is also supported by the Chan Zuckerberg Biohub and NIH GM102365. A.G. is supported by the National Institutes of Health for the Pharmacogene Variation Consortium (R24GM123930). P.S.T is supported by 1HL-101388 (NIH-NHLBI). We thank Kenneth Thummel at the University of Washington and Erin Schuetz at St. Jude Children’s Research Hospital for providing the liver bank resources to conduct this work.

## Declaration of Interests

RBA is a stockholder in Personalis.com, 23andme.com, and Youscript.com.

