## Supplemental Data for "Transfer learning enables prediction of *CYP2D6* haplotype function"

Supplementary Table 1. *CYP2D6* Star allele function predictions

| ***CYP2D6* Star Allele** | **Curated Function** | **Hubble Predicted Function** |
| --- | --- | --- |
| *1 | Normal | Normal |
| *2 | Normal | Normal |
| *3 | No function | No function |
| *4 | No function | No function |
| *6 | No function | No function |
| *7 | No function | No function |
| *8 | No function | No function |
| *9 | Decreased function | Decreased function |
| *10 | Decreased function | Decreased function |
| *11 | No function | No function |
| *12 | No function | No function |
| *14 | Decreased function | No function |
| *14 | Decreased function | No function |
| *15 | No function | No function |
| *17 | Decreased function | Decreased function |
| *18 | No function | No function |
| *19 | No function | No function |
| *20 | No function | No function |
| *21 | No function | No function |
| *22 | Uncurated | Decreased function |
| *23 | Uncurated | Normal |
| *24 | Uncurated | Normal |
| *25 | Uncurated | Normal |
| *26 | Uncurated | Normal |
| *27 | Normal | Normal |
| *28 | Uncurated | Decreased function |
| *29 | Decreased function | Decreased function |
| *30 | Uncurated | Decreased function |
| *31 | No function | No function |
| *32 | Uncurated | Decreased function |
| *33 | Normal | Normal |
| *34 | Normal | Normal |
| *35 | Normal | Normal |
| *36 | No function | No function |
| *37 | Uncurated | Decreased function |
| *38 | No function | No function |
| *39 | Normal | Normal |
| *40 | No function | No function |
| *41 | Decreased function | Decreased function |
| *42 | No function | No function |
| *43 | Uncurated | Normal |
| *44 | No function | No function |
| *45 | Normal | Normal |
| *46 | Normal | Normal |
| *47 | No function | No function |
| *48 | Normal | Normal |
| *49 | Decreased function | Decreased function |
| *50 | Decreased function | Decreased function |
| *51 | No function | No function |
| *52 | Uncurated | Decreased function |
| *53 | Normal | Normal |
| *54 | Decreased function | Decreased function |
| *55 | Decreased function | Decreased function |
| *56 | No function | No function |
| *57 | No function | No function |
| *58 | Uncurated | Decreased function |
| *59 | Decreased function | Decreased function |
| *60 | No function | No function |
| *62 | No function | No function |
| *64 | Uncurated | No function |
| *65 | Uncurated | Decreased function |
| *69 | No function | No function |
| *70 | Uncurated | Decreased function |
| *71 | Uncurated | Decreased function |
| *72 | Decreased function | No function |
| *73 | Uncurated | Decreased function |
| *74 | Uncurated | Normal |
| *75 | Uncurated | No function |
| *81 | Uncurated | Decreased function |
| *82 | Uncurated | Normal |
| *83 | Uncurated | Normal |
| *84 | Decreased function | Decreased function |
| *85 | Uncurated | Normal |
| *86 | Uncurated | Normal |
| *87 | Uncurated | Decreased function |
| *88 | Uncurated | Decreased function |
| *89 | Uncurated | Normal |
| *90 | Uncurated | Normal |
| *91 | Uncurated | Decreased function |
| *92 | No function | No function |
| *93 | Uncurated | Normal |
| *94 | Uncurated | Decreased function |
| *95 | Uncurated | Decreased function |
| *96 | No function | No function |
| *97 | Uncurated | Normal |
| *98 | Uncurated | Decreased function |
| *99 | No function | No function |
| *100 | No function | No function |
| *101 | No function | No function |
| *102 | Uncurated | Decreased function |
| *103 | Uncurated | Decreased function |
| *104 | Uncurated | Decreased function |
| *105 | Uncurated | Decreased function |
| *106 | Uncurated | Normal |
| *107 | Uncurated | Normal |
| *108 | Uncurated | Normal |
| *109 | Uncurated | Decreased function |
| *110 | Uncurated | Decreased function |
| *111 | Uncurated | Decreased function |
| *112 | Uncurated | Normal |
| *113 | Uncurated | Decreased function |
| *114 | No function | No function |
| *115 | Uncurated | Decreased function |
| *116 | Uncurated | Normal |
| *117 | Uncurated | Decreased function |
| *118 | Uncurated | Normal |
| *119 | Uncurated | Decreased function |
| *120 | Uncurated | No function |
| *121 | Uncurated | Decreased function |
| *122 | Uncurated | Normal |
| *123 | Uncurated | Decreased function |
| *124 | Uncurated | Decreased function |
| *125 | Uncurated | Normal |
| *126 | Uncurated | Normal |
| *127 | Uncurated | Decreased function |
| *128 | Uncurated | Decreased function |
| *129 | Uncurated | Decreased function |
| *130 | Uncurated | Normal |
| *131 | Uncurated | Normal |
| *132 | Uncurated | No function |
| *133 | Uncurated | Decreased function |
| *134 | Uncurated | Normal |
| *135 | Uncurated | Normal |
| *136 | Uncurated | Normal |
| *137 | Uncurated | Normal |
| *138 | Uncurated | No function |
| *139 | Uncurated | Normal |

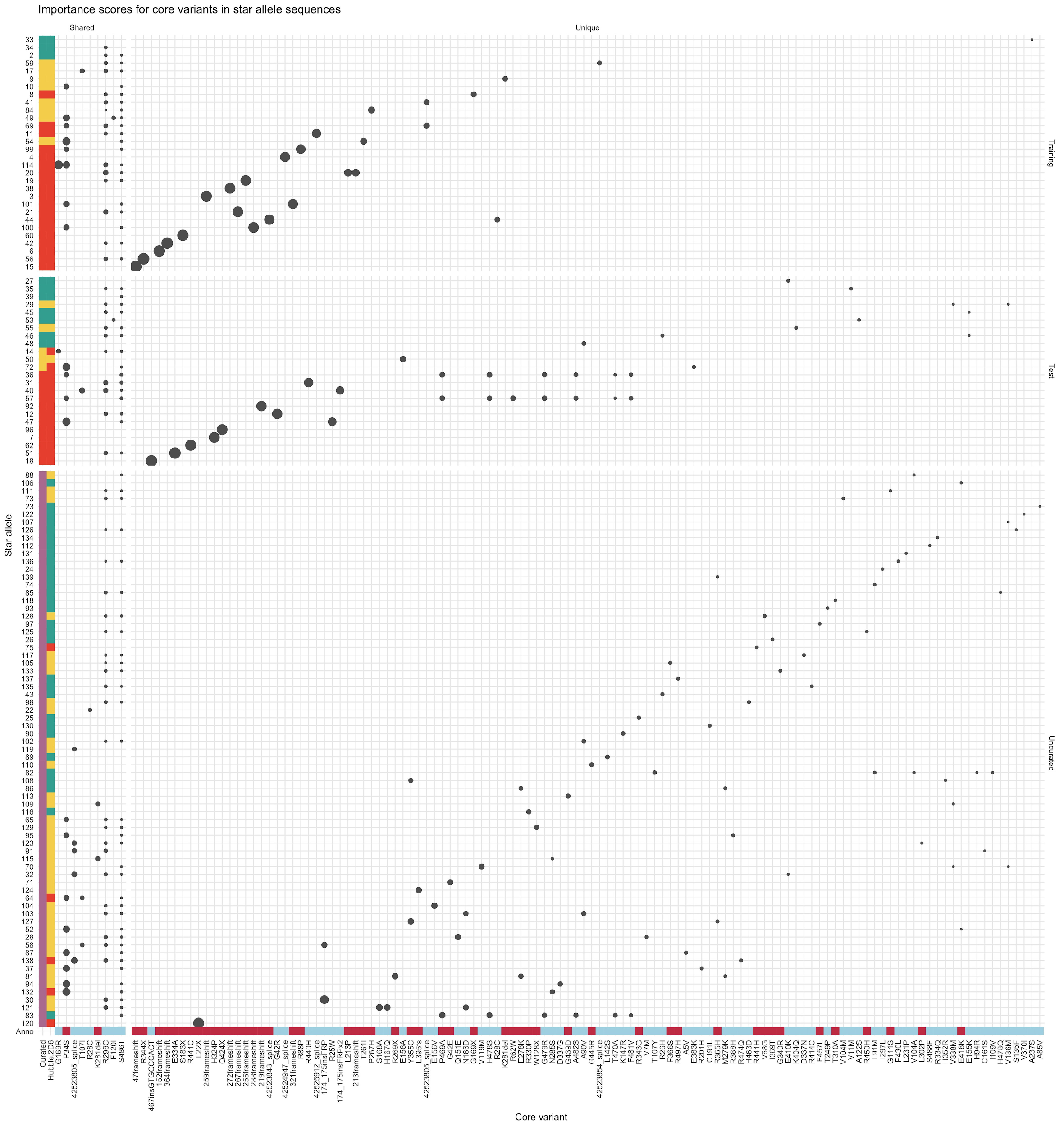

Supplementary Figure 1. Importance scores for core variants in each star allele used for training and test of Hubble.2D6, as well as uncurated star alleles. Star alleles are along the y-axis and core variants (both amino acid changes and non-coding changes) are listed along the x-axis. Each dot represents the importance of the core variant to the final prediction as determined by DeepLIFT. The size of the dot represents the value of the importance score, with larger dots indicating variants with larger importance scores, typically associated with a negative impact on function. Star alleles are annotated with the curated function as well as the Hubble.2D6 predicted function. Star alleles are divided along the y-axis between star alleles that were included in the training data (top) and those used as test samples (bottom). Star alleles are sorted by the sum of the importance scores, with those with the largest sums at the bottom. Core variants are divided along the x-axis by those that are uniquely in either the training or test samples (right), and those that are shared between star alleles in train and test (left). Core variants are sorted by their mean importance score across all star alleles. Core variants are annotated with the deleteriousness annotation used in the functional variant representation.
